# CARACAS, a novel automated tool for Cardiac Artifact Removal in Absence of CArdiac Signal

**DOI:** 10.1101/2025.09.02.673728

**Authors:** Pierre Champetier, Delphine Oudiette, Thomas Andrillon, Maximillien Chaumon

## Abstract

**Background:** EEG recordings can contain cardiac related artifacts. Independent Component Analysis (ICA) followed by removal of cardiac Independent Components (ICs) is a powerful and widely used strategy for artifact correction. Most existing methods for automatic labeling of cardiac ICs require a simultaneously recorded ECG (e.g., to compute correlation with the IC time course). However, ECG is not always available. To address this limitation, we developed CARACAS (Cardiac Artifact Removal in Absence of CArdiac Signal), a novel tool that identifies cardiac ICs using only the IC time courses.

**New method:** Because cardiac ICs exhibit temporal profiles highly similar to ECG signals, we used an existing tool designed to detect cardiac events (R waves) in ECG signals and applied it to each IC time course. Analysis of the detected events enabled the differentiation of cardiac ICs from non-cardiac ICs, where unrelated signal variations are incorrectly identified as cardiac events. Using the 375 EEG-ECG recordings of the open-source dataset OpenNeuro ds003690, we compared the performances of three algorithms: CARACAS, IClabel (a generic IC classifier which does not require ECG), and correlation with ECG channel.

**Results (comparison with existing methods):** A total of 21,375 ICs were manually and automatically classified. CARACAS achieved high performance (sensitivity = 0.960, specificity = 0.976), substantially outperforming ICLabel (sensitivity = 0.210, specificity = 0.999) and approaching the performance of ECG correlation method (sensitivity = 0.975, specificity = 0.998).

**Conclusion:** We present a reliable ECG-free algorithm for cardiac IC detection in EEG. CARACAS provides a practical solution when ECG is unavailable, and is implemented in the SASICA toolbox.

**Highlights:**

- Cardiac independent component (IC) removal after ICA corrects cardiac EEG artifacts.
- Most automatic cardiac IC detectors require a simultaneously recorded ECG.
- We developed CARACAS, a novel ECG-free method for automatic cardiac IC labeling.
- CARACAS achieved a sensitivity of 0.960 and a specificity of 0.976 on 21,375 ICs.
- CARACAS outperforms ICLabel, and is available in SASICA toolbox (command line & GUI).

**Graphical abstract:** 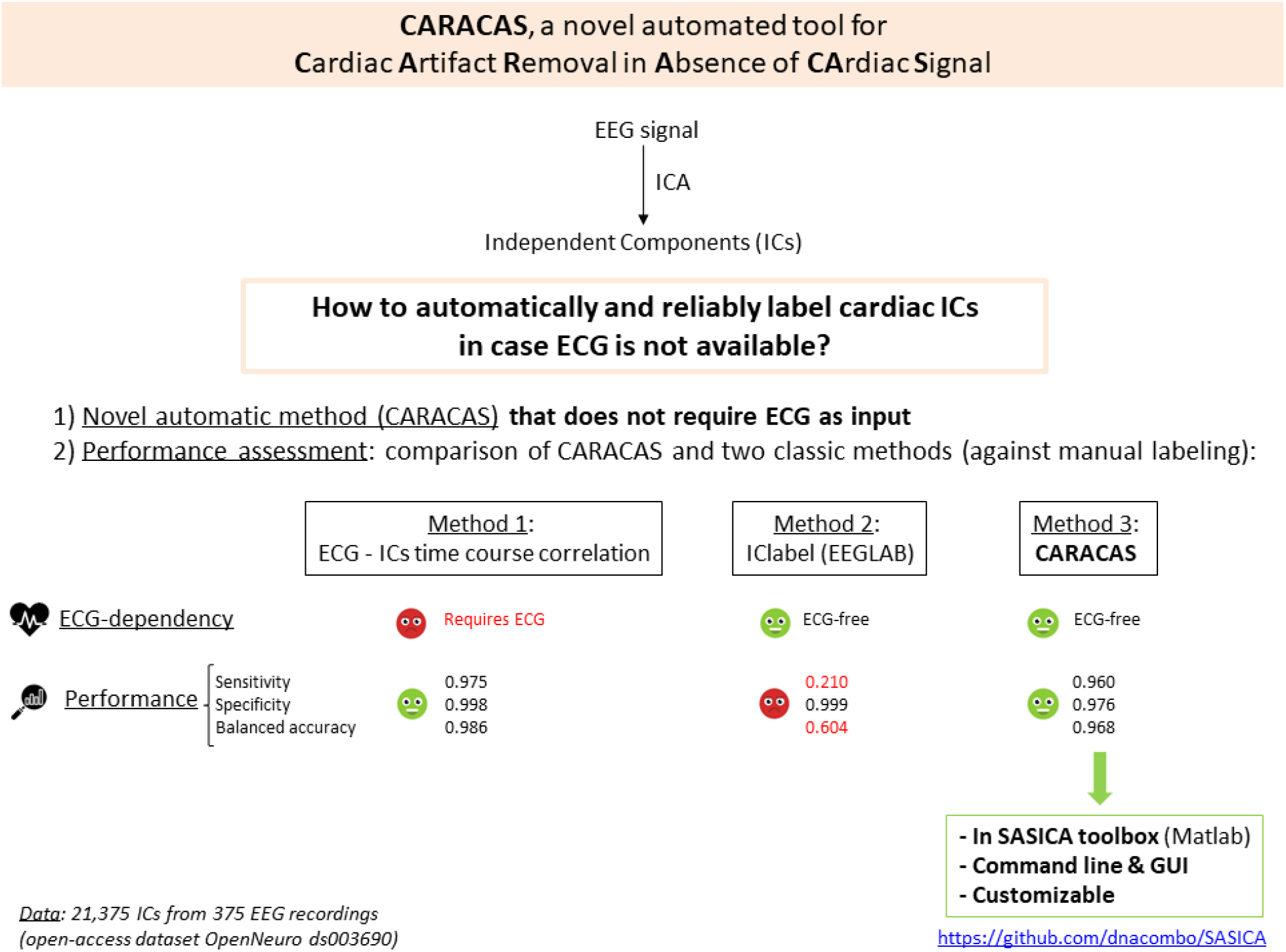

## Introduction

Electroencephalography (EEG) is a widely used technique in both research and clinical settings for recording brain activity. Its affordability, non-invasiveness, and ease of use make it a valuable tool in neuroscience. However, EEG signals are often contaminated by various artifacts, both biological (e.g., eye movements, cardiac activity) and non-biological (e.g., line noise), which can significantly compromise data quality and interpretation. Independent Component Analysis (ICA) is one of the most common techniques used to isolate and remove such artifacts (Urigüen and Garcia-Zapirain, 2015). By decomposing the EEG signal into temporally independent components (ICs), ICA allows the identification and exclusion of components associated with artifacts (e.g., blinks, saccades, cardiac activity) before reconstructing a cleaned EEG signal. While this identification can be performed manually, the process is time-consuming, subjective, and not scalable to the large datasets that are increasingly common in modern neuroscience. To address this challenge, automated IC classification methods have emerged as essential tools. Cardiac-related ICs, in particular, can be reliably identified when a simultaneous electrocardiogram (ECG) is available, by correlating IC time courses with the ECG signal, as implemented in the SASICA toolbox (Chaumon et al., 2015). While ECG recording is standard in sleep recordings (polysomnography), it is less systematic in wake or routine clinical EEG protocols, and is not included in EEG recommendations (Niso et al., 2022; Pernet et al., 2020). Furthermore, the growing use of portable multi-channel EEG systems for ecological recordings (at home or during daily activities) often comes with a deliberate reduction of sensors to improve ergonomics, making ECG unavailable in many such setups. Thus, automatic labeling of cardiac ICs remains a challenge in datasets lacking simultaneous ECG. While generic classifiers such as ICLabel (integrated into EEGLAB) offer automated solutions (Pion-Tonachini et al., 2019), they are not specifically optimized for cardiac components and can show limited performance in this regard. More recent methods have attempted to fill this gap by leveraging alternative strategies using rule-based classifiers (Issa et al., 2019; Tamburro et al., 2019) or neural network models (Arnau et al., 2023). Yet, these tools lack robust validation, are not always open-access and lack a user-friendly interface, hindering widespread adoption. In this context, we present CARACAS (Cardiac Artefact Removal in Absence of CArdiac Signal), a new open-source tool specifically designed to automatically identify cardiac-related ICs without the need for an ECG signal. Our goal is to provide an accessible, reliable, and efficient solution to improve artifact rejection workflows in EEG processing pipelines.

## Methods

### EEG datasets

CARACAS was tested on the open-access EEG dataset OpenNeuro ds003690 (doi:10.18112/openneuro.ds003690.v1.0.0). This dataset includes n = 375 recordings (from 36 young adults and 39 older adults). For each subject, 5 recordings were available: one during passive task (4-min passive listening of auditory stimulus) and four during active tasks (two runs during a go/no go task and two runs during a passive simple reaction time task, each 8-min). Each recording comprised signals from 64 EEG electrodes and one ECG lead, stored in EEGLAB format. Data collection procedure was detailed in (Ribeiro and Castelo-Branco, 2019). Briefly, EEG signals were acquired using a 64-channel Neuroscan system at a sampling frequency of 500 Hz. Scalp electrodes were positioned according to the International 10-20 system. To minimize motion artifacts, participants’ heads were stabilized using a chin and forehead rest during recording.

### Preprocessing

Signal processing was performed using the FieldTrip toolbox (fetched on 2025/04/23 from https://github.com/fieldtrip/fieldtrip) (Oostenveld et al., 2011). Two mastoid electrodes M1 and M2 were removed and the remaining electrodes were re-referenced to the average. A finite impulse response (FIR) bandpass filter between 0.1 and 35 Hz was applied with default parameters of the FieldTrip version used. The EEG signal was time-locked to “cue” (see details in (Ribeiro and Castelo-Branco, 2019)) with a pre-stimulus time of 0.2 seconds and post-stimulus time of 0.5 seconds. Pre-stimulus baseline was subtracted. ICA was applied using the runica method with default parameter values on EEG channels, yielding 57 ICs per recording. No other preprocessing was applied.

### Labeling of cardiac IC using pre-existing procedures

#### Manual IC labeling

The 21,375 ICs (375 recordings x 57 ICs) were manually labeled as “cardiac” or “non-cardiac” by an experienced EEG researcher based on visual inspection of their time courses and topographies. ICs were classified as cardiac only when they exhibited a clear, high-amplitude, and regular ECG-like pattern.

#### Automatic IC labeling

ICs were also automatically classified as “cardiac” or “non-cardiac” using two established algorithms.

1. The IClabel function (Pion-Tonachini et al., 2019) of EEGLAB toolbox (MATLAB) was applied on ICA outputs. For each IC, IClabel returns the probability of originating from one of seven sources (brain, muscle, eye, heart, line noise, channel noise, or other). Following the authors’ recommendations, an IC was labeled as cardiac when “heart” was the category with the highest probability.
2. The “correlation with channel” method of the SASICA toolbox (MATLAB) was used to compute the correlation between each IC time course and the simultaneously recorded ECG. An IC was labeled as cardiac when this correlation exceeded 0.6 (default threshold in the toolbox). Hereafter, this labeling procedure is referred to as CORR.

### Labeling of cardiac IC using CARACAS

CARACAS algorithm analyzes each IC-time course separately. The key steps of the procedure to identify cardiac ICs with CARACAS are summarized in **Figure 1**.

**Fig. 1:**
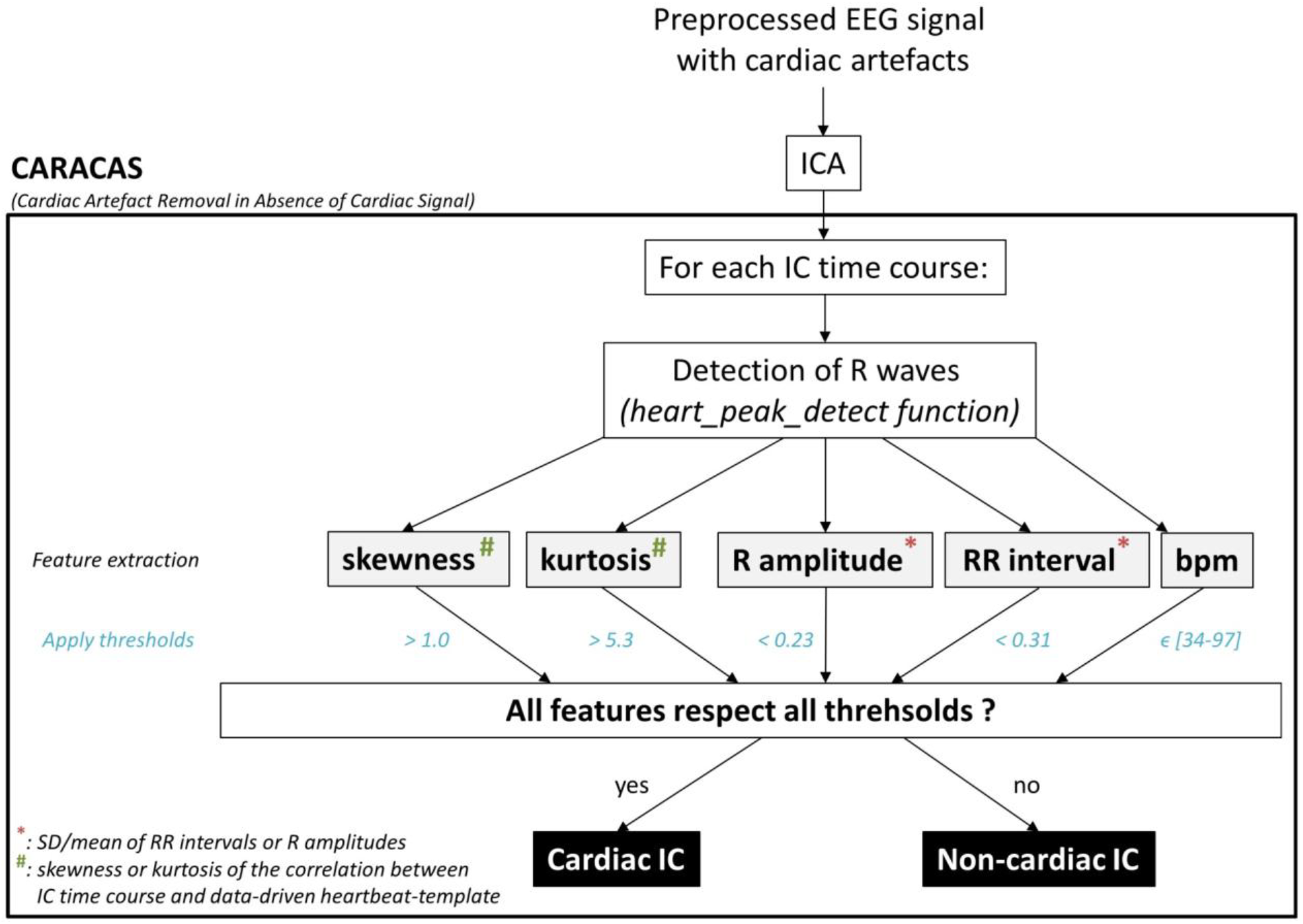
Flowchart of CARACAS key steps to identify cardiac ICs. Following an ICA, CARACAS uses the IC time courses as inputs. In each time course, it detects cardiac events using an existing function that uses relative thresholds. Then, it extracts 5 features grasping complementary aspects of the IC time course. Each feature value is compared to a predefined threshold, allowing labeling of the IC as “cardiac” or “non-cardiac”. <u>Abbreviations:</u> bpm (beat per minute), ICA (Independent Component Analysis), IC (Independent Component).

#### 1 R labeling

In the first step, an existing algorithm (GitHub repository: https://github.com/dnacombo/heart_functions, previously used in (Buot et al., 2021)) originally developed to label cardiac events (PQRST) in ECG recordings was applied without modification to each IC time course. Thus, this function examines each IC in turn as if it were an ECG channel (hereafter called “candidate ECG”) and implements a two-stage algorithm for comprehensive cardiac peak detection in the IC time course. Briefly, this algorithm works as follows: first, the candidate ECG signal is preprocessed with bandpass filtering (default: 1-100 Hz), followed by z-score normalization and squaring to enhance R-wave prominence. Most prominent R-peak candidates are identified using an amplitude threshold (default: z=10) applied to the squared signal with minimum inter-beat interval constraints (default: 0.35s). Due to the relative nature of the threshold, R-peak labels are nearly always assigned, even in non-cardiac ICs. A representative heartbeat template is constructed by averaging signal segments around the detected R-peaks (±0.5s windows). This template is then convolved with the entire filtered candidate ECG signal to generate a correlation time series, from which the remaining R-peaks are identified as correlation maxima exceeding a threshold (default: 0.6). Subsequently, the algorithm detects the remaining PQRST complex components relative to each R-peak: Q-waves as minima within 50ms preceding R-peaks, P-waves as maxima within 250ms preceding Q-waves, S-waves as minima within 100ms following R-peaks, and T-waves as maxima between S-waves and a maximum QT interval of 420ms. This procedure always labels cardiac events, in a fixed PQRST sequence, even in non-cardiac ICs. Only R-peaks are used in CARACAS.

#### 2 Feature extraction

The second step involves extracting features to distinguish between cardiac and non-cardiac ICs. Five features are extracted (hereafter 1 to 5). (1) The skewness and (2) the kurtosis of the distribution of correlations between the heartbeat-template and the IC-time course are extracted. Besides, the standard deviation/mean ratio (SD/mean) is computed for the (3) R amplitudes and the (4) RR intervals, using the detected R peaks on each IC-time course. To avoid biases due to R detection errors (false positive or misses) in true cardiac ICs, the 30% biggest values of all RR intervals and the 30% most extreme values of R amplitudes are excluded from each distribution before computing the SD/mean ratios. (i.e., only the 0th–70th percentile for RR intervals and the 15th–85th percentiles for R amplitudes are kept). Finally, the (5) beat per minute (bpm) is calculated, based on the detected R peaks on each IC-time course. Together, these features grasp complementary aspects of the IC-time course. The skewness and kurtosis of the correlation with heartbeat-template give insight into the pattern consistency of detected cardiac events. The SD/mean ratios assess the regularity in time and amplitude of detected R waves (i.e., the main cardiac event). Finally, the bpm allows checking if the overall temporal density of detected cardiac events is physiologic.

#### 3 Comparison to thresholds

In the third step, each of the five extracted features is compared against predefined thresholds. An IC is labeled as cardiac if it simultaneously meets all of the following criteria: (1) skewness > 1.0, (2) kurtosis > 5.3, (3) SD/mean(R amplitudes) < 0.23, (4) SD/mean(RR intervals) < 0.31, and (5) bpm ∈ [34-97/min].

These thresholds, which remain easily customizable by the user, were first approximated by visual inspection of the distributions of each feature for manually labeled cardiac and non-cardiac ICs. Approximate thresholds were visually selected at the tails of the cardiac IC distribution, in order to exclude non-cardiac ICs while retaining nearly all cardiac ICs. This yielded the following initial values: skewness > 1.0, kurtosis > 5.0, SD/mean(R amplitudes) < 0.25, SD/mean(RR intervals) < 0.33, and bpm ∈ [35-100/min].

Subsequently, these approximated thresholds were refined and validated using a nested 5-fold cross-validation procedure, to obtain an unbiased estimate of the model’s generalization performance and to reduce the risk of overfitting during model selection. In the outer loop, data was divided into five folds. In each iteration, 80% of the data were used for optimization (training) and the remaining 20% were held out for final performance evaluation. Within each outer training set, an inner 5-fold cross-validation was performed to optimize the threshold values. For this inner loop, the fminsearch function in MATLAB iteratively adjusted the thresholds—starting from random initial values within ±10% of the visually defined ranges—to maximize classification performance, quantified by balanced accuracy (see Performance Evaluation). Averages of the optimal thresholds from the inner loop were then applied to the corresponding outer test set to assess performance. This process was repeated across the five outer folds, ensuring that parameter tuning and performance evaluation were carried out on independent subsets of data. The final default thresholds implemented in CARACAS correspond to the median values obtained across the outer folds.

The output of the CARACAS algorithm provides the user with: (1) a list of ICs identified as cardiac, and (2) for each component, the measures defined above, and which ones passed the set threshold and were therefore classified as not cardiac. The algorithm has been implemented in the SASICA toolbox (version 1.4 and later, https://github.com/dnacombo/SASICA), allowing visual inspection of each IC, including CARACAS outputs (**Figure 2**).. The code used for the analyses in this paper can be found here: https://github.com/dnacombo/CARACASpaper.git.

**Fig. 2:**
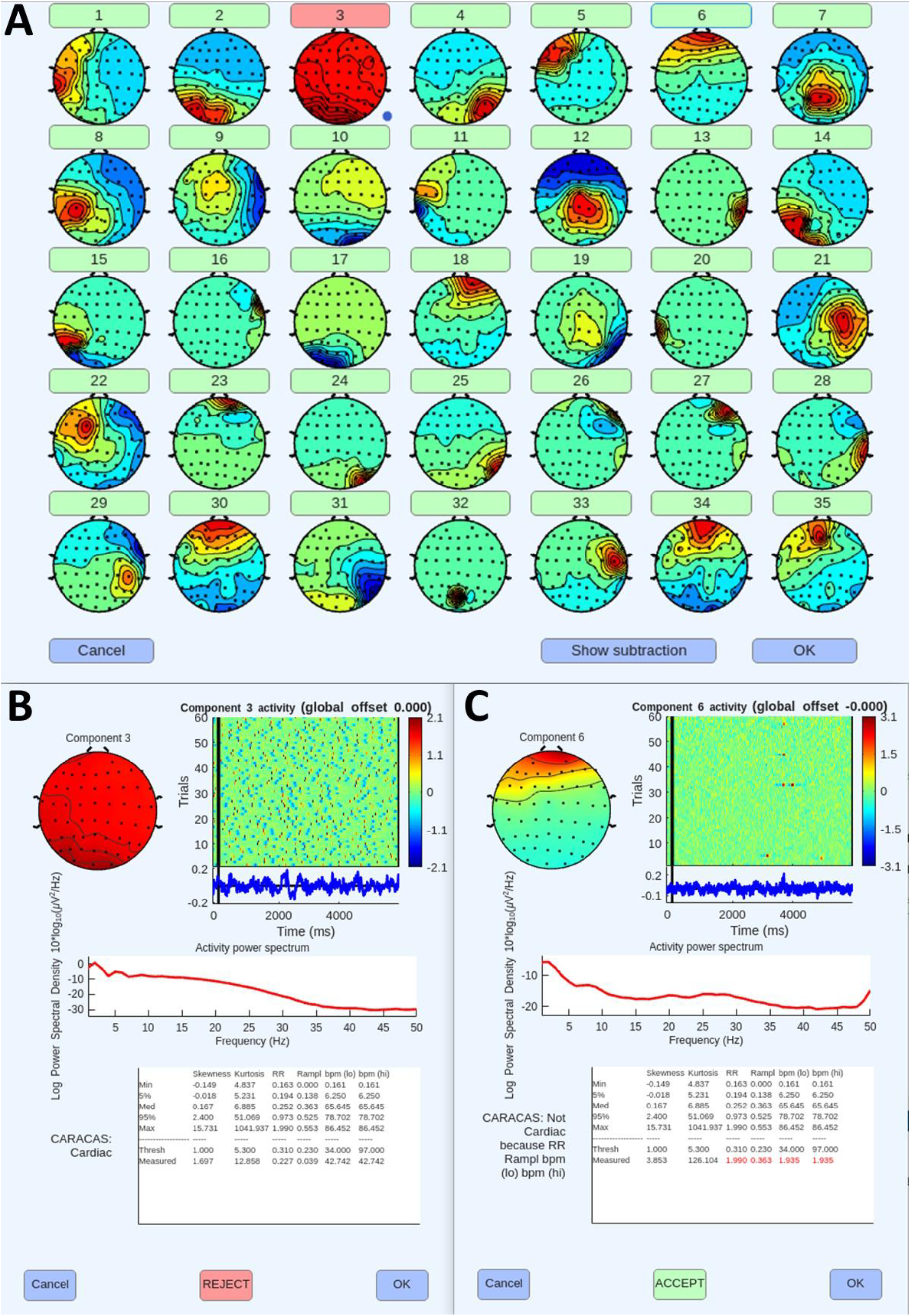
Graphical user interface of CARACAS outputs within the SASICA toolbox. (A) SASICA window displaying all IC topographies (35 ICs in this example). IC numbers are indicated on buttons colored green or red, with red indicating ICs labeled as cardiac by CARACAS (e.g., IC #3). (B-C) Clicking on a button displays detailed features of the selected IC: (B) example of a cardiac IC; (C) example of a blink IC. For each IC, the following information is shown: topography, activity for each trial (ERP image), average time course across trials, power spectrum, and CARACAS metrics. For each CARACAS metric (skewness, kurtosis, R-R interval, R amplitudes, and bpm range), the panel displays the IC’s measured value, the classification threshold, and descriptive statistics across all ICs in the recording (minimum, 5^th^ percentile, median, 95^th^ percentile, and maximum). For ICs labeled as non-cardiac, metrics not meeting the threshold are highlighted in red.

### Performance evaluation

To assess the CARACAS performances, we compared the labels of each IC (cardiac or non-cardiac) attributed by the algorithm to those attributed manually. Sensitivity (i.e., true positive rate) and specificity (i.e., true negative rate) were calculated as follows: 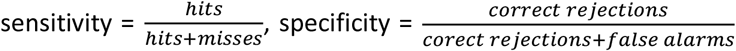. Balanced accuracy 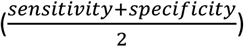 was also computed to assess the overall performance, while taking into account the imbalance of the labels of the IC (see Results section). The same performance metrics were computed for IClabel and CORR using the same procedure. To illustrate the variability and stability of the nested cross-validation results used for CARACAS, performance metrics obtained for each outer fold on their respective test set are provided in the **Supplementary Table 1**.

## Results

From each of the 375 recordings, ICA procedure yielded in 57 ICs. Manual inspection identified one cardiac IC in most recordings, resulting in a total of 308 cardiac ICs (1.44% of the 21,375 ICs).

The three automated algorithms demonstrated varying performance (**Table 1**). Among the 308 manually identified cardiac ICs, only 65 were correctly classified by ICLabel (sensitivity = 0.210), compared with 296 by CARACAS (sensitivity = 0.960) and 298 by CORR (sensitivity = 0.975). False positives were rare across all three algorithms, leading to specificity consistently greater than 0.98. Consequently, ICLabel showed the weakest overall performance (balanced accuracy = 0.604), whereas CARACAS (balanced accuracy = 0.968) closely approached the performance of CORR (balanced accuracy = 0.986).

**Table 1.**
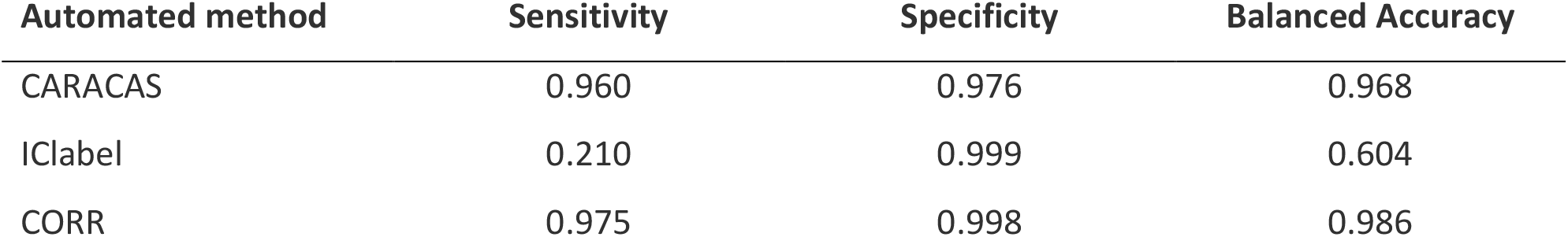
Performances of automated cardiac classifiers when considering all ICs (n= 21,375)

| Automated method | Sensitivity | Specificity | Balanced Accuracy |
| --- | --- | --- | --- |
| CARACAS | 0.960 | 0.976 | 0.968 |
| ICLabel | 0.210 | 0.999 | 0.604 |
| CORR | 0.975 | 0.998 | 0.986 |

We repeated the analysis restricting it to the 10 most important ICs per recording (i.e., the first 10 components; n = 3,750), as these components are most likely to contain substantial EEG artifacts. Results remained highly consistent with the full analysis, with CARACAS achieving improved sensitivity (0.965) and balanced accuracy (0.970), nearly matching CORR’s balanced accuracy (0.983, **Table 2**).

**Table 2.** Performances of automated cardiac classifiers when considering only the first 10 ICs of each recording (n= 3,750)

| Automated method | Sensitivity | Specificity | Balanced Accuracy |
| --- | --- | --- | --- |
| CARACAS | 0.965 | 0.974 | 0.970 |
| IClabel | 0.211 | 0.997 | 0.604 |
| CORR | 0.972 | 0.995 | 0.983 |

On the contrary, CARACAS performance was lower on the two last ICs (i.e., those explaining the lowest variance in the EEG signal). Manual inspection showed none of these were cardiac-related, but the algorithm incorrectly labeled some of them as cardiac (specificity restricted to those last two components = 0.803).

Finally, we repeated the main analysis when applying stricter high-pass filtering (at 1 Hz) to EEG data prior to ICA, a step sometimes recommended to improve ICA decomposition (Klug and Gramann, 2021). Similar results were obtained, with CARACAS performances clearly outperforming IClabel ones (**Supplementary Table 2**).

## Discussion

In this study, we introduced CARACAS, a novel open-source tool for the automatic identification of cardiac ICs in EEG data without requiring a simultaneously recorded ECG. Across 375 recordings, CARACAS demonstrated high sensitivity and specificity, substantially outperforming ICLabel, a widely used generic classifier that does not require ECG, and approaching the performance of CORR, the classic algorithm that relies on ECG recordings. CARACAS therefore provides an efficient ECG-free solution for the automatic labeling of cardiac ICs, a critical step in EEG artifact removal. Such an approach is particularly relevant when ECG is unavailable due to technical issues, deliberately omitted to reduce setup complexity (e.g., in ecological or multimodal settings), degraded, or simply absent from archival and large-scale datasets with heterogeneous recording protocols.

Previous ECG-free approaches specifically developed to detect cardiac ICs have been limited in scope and validation. For example, the method proposed by Issa et al. was tested in only seven resting-state recordings (Issa et al., 2019), while the algorithm by Tamburro et al. was evaluated using accuracy (Tamburro et al., 2019), a maladapted metric. Accuracy, defined as (true positives + true negatives) / total, can be misleading for strongly unbalanced datasets, as cardiac ICs represent only a small fraction of all the ICs (∼1% in our case). For instance, in a dataset of 10,000 ICs with 1% cardiac ICs, accuracy of an algorithm that classifies all ICs as non-cardiac would be (0 + 9,900) / 10,000 = 0.99. Although this value appears excellent, it is entirely driven by the high specificity: since 99% of ICs are non-cardiac, the algorithm achieves near-perfect performance on the majority class, despite never detecting a single cardiac IC (sensitivity = 0). By contrast, balanced accuracy, which equally weights sensitivity and specificity, more accurately reflects chance-level performance: (0 + 1) / 2 = 0.5. Unlike previous approaches, which were limited in scope and relied on inadequate validation metrics, CARACAS was validated on a large open-access dataset and evaluated using balanced accuracy, providing a more realistic assessment of classification performance in imbalanced conditions.

CARACAS builds on the observation that cardiac ICs exhibit time courses closely resembling ECG signals. The method detects putative cardiac events in all IC time courses, extracts five complementary features from these events, and compares each of these features to thresholds. The multi-criteria design, in which an IC is discarded if it fails to meet even one criterion, effectively reduces the likelihood of false negatives. For example, time course of blink-related ICs also exhibit stereotypical repetitive structure, but fail to meet other criteria such as physiological density of events or temporal regularity, thereby avoiding misclassification.

CARACAS has several strengths.

1. It does not require a simultaneously recorded ECG, which makes it applicable to datasets where ECG is missing or of poor quality.
2. It enables reliable and automatic IC classification, making it scalable to large datasets while eliminating the subjectivity of manual inspection.
3. It has been validated on an open-access dataset of 375 recordings, including both resting-state and several task conditions. Evaluation was performed using balanced accuracy, which accounts for the strong class imbalance between cardiac and non-cardiac ICs. CARACAS proved highly efficient, with performance comparable to CORR, one of the gold-standard procedures requiring ECG. Importantly, its sensitivity further improved when focusing on ICs explaining the largest portion of variance, those most likely to introduce substantial artifacts into EEG data. Across all analyses, CARACAS maintained a very low false-positive rate, with specificity exceeding 0.98 when all ICs were considered.
4. It labels each IC independently. Thus, its performance does not depend on the number of ICs. This allows for reliable use across a wide range of decompositions, as demonstrated in our analysis restricted to the first 10 ICs.
5. It provides several practical advantages: it has low computation cost, allows flexible adjustment of thresholds, and is fully open-source within the SASICA toolbox.

However, some limitations must be acknowledged. First, CARACAS relies on empirically optimized thresholds for feature selection. Although these were refined through cross-validation, they may require adaptation for datasets with different recording conditions, populations, or noise profiles. Second, validation was performed on a single dataset collected during wakefulness. Although the dataset includes several types of recordings (resting-state, active or passive tasks), broader evaluation on diverse EEG systems and paradigms (e.g., sleep studies, mobile EEG during naturalistic tasks, clinical populations) will be necessary to assess generalizability.

In conclusion, CARACAS provides the community with a transparent, efficient, and ECG-free tool for the labeling of cardiac ICs in ICA-decomposed EEG data. Its implementation in the SASICA toolbox ensures straightforward integration into existing preprocessing pipelines, thereby lowering the barrier for large-scale, reproducible artifact rejection. Future developments will include the creation of a graphical user interface (GUI) to facilitate adoption by non-programmers. CARACAS thus represents a reliable and practical solution for EEG datasets in which ECG is unavailable or of insufficient quality.

## Supplementary material

**Supplementary Table 1.** Optimized threshold values and corresponding performance metrics for each outer fold of the nested cross-validation procedure.

| Outer fold | Sensitivity | Specificity | Balanced Accuracy |
| --- | --- | --- | --- |
| 1 | 0.951 | 0.977 | 0.964 |
| 2 | 0.959 | 0.978 | 0.968 |
| 3 | 0.859 | 0.979 | 0.919 |
| 4 | 0.982 | 0.980 | 0.981 |
| 5 | 0.911 | 0.985 | 0.948 |

**Supplementary Table 2.** Performances of automated cardiac classifiers with a 1-Hz high-pass filter on EEG data prior to ICA.

| Automated method | Sensitivity | Specificity | Balanced Accuracy |
| --- | --- | --- | --- |
| CARACAS | 0.842 | 0.970 | 0.906 |
| IClabel | 0.337 | 0.999 | 0.668 |
| CORR | 0.959 | 1 | 0.979 |

*All ICs were used in this analysis (n= 21,375).*

